# Intranasal immunization with CPAF combined with cyclic-di-AMP induces a memory CD4 T cell response and reduces bacterial burden following intravaginal infection with *Chlamydia muridarum*

**DOI:** 10.1101/2024.09.20.614154

**Authors:** Taylor B. Poston, Jenna Girardi, Marie Kim, Peter Zwarycz, A. Grace Polson, Kacy Yount, Courtne Hanlan, Ian Jaras Salas, Sarah Mae Lammert, Daisy Arroyo, Tony Bruno, Manhong Wu, James Rozzelle, Jeff Fairman, Aaron Esser-Kahn, Toni Darville

## Abstract

*Chlamydia trachomatis* (Ct) is the most common bacterial sexually transmitted infection globally, and a vaccine is urgently needed to stop transmission and disease. Chlamydial Protease Activity Factor (CPAF) is an immunoprevalent and immunodominant antigen for CD4 T cells and B cells, which makes it a strong vaccine candidate. Due to the tolerogenic nature of the female genital tract (FGT) and its lack of secondary lymphoid tissue, effective induction of protective cell-mediated immunity will likely require potent and safe mucosal adjuvants. To address this need, we produced CPAF in a cell-free protein synthesis platform and adjuvanted it with the TLR9-agonist CpG1826, STING (stimulator of interferon genes) agonist cyclic-di-AMP (CDA), and/or the squalene oil-in-water nanoemulsion, AddaS03. We determined that intranasal immunization with CPAF plus CDA was well tolerated in female mice, induced CD4 T cells that produced IL-17A or IFNγ, significantly reduced bacterial shedding, and shortened the duration of infection in mice intravaginally challenged with *Chlamydia muridarum*. These data demonstrate the potential for CDA as a mucosal adjuvant for vaccines against Chlamydia genital tract infection.

## INTRODUCTION

Genital *Chlamydia trachomatis* (CT) infection and its associated diseases represent a significant global health burden, with ∼131 million new cases occurring annually1. While antibiotic treatment is highly effective, more than 70% of infections are asymptomatic, leading to many cases going undetected and untreated^2, 3^. When infection ascends from the cervix to the endometrium and oviducts, it induces tissue-damaging inflammation, which can result in pelvic inflammatory disease (PID) and potential infertility. Despite existing screening programs, infection rates continue to rise^4^. This underscores the urgent need for a vaccine that combats this ongoing epidemic and associated complications.

We previously evaluated antibody and T cell responses in CT-exposed women enrolled in the T cell Response Against Chlamydia (TRAC) study^5, 6^. Whole immunoproteome antibody profiling of sera obtained from 222 TRAC participants revealed that CPAF (CT858) was the most immunogenic protein, recognized by 75% of participants, and had the highest average antibody levels among all antigens^7^. Additionally, we recently determined that CPAF was also the most immunoprevalent and immunodominant CD4 T cell antigen screened in this cohort^8^. These data demonstrate the potential of CPAF as a promising candidate for a human vaccine antigen.

Most vaccines against bacterial pathogens induce antibody as a primary mode of protection. However, chlamydial protection is dependent on CD4 T cell-mediated responses^9–12^. Until the past decade, the development of vaccines against intracellular bacterial pathogens, which require cell-mediated immunity for protection, was hindered by the absence of safe and effective adjuvants that could induce Th1 and Th17 responses^13^. Early studies using CPAF as an immunogen demonstrated protection when CPAF was adjuvanted with IL-12 or multiple doses of CpG before and after CPAF immunization^14, 15^. However, the toxicity of IL-12 administration and the implausibility of delivering CpG on consecutive days make these approaches unsuitable for human vaccination. Bacterial agonists such as c-di-AMP (CDA) show promise as adjuvants with a good safety record for mice and humans^16–18^. CDA activates the cytosolic STING pathway that elicits inflammatory cytokines and interferons to drive robust Th1/17 responses^19–22^. CpG and AS03 have shown effectiveness in eliciting CD4 T cell responses in humans^23–25^.

Due to the good safety and immunogenicity profiles of CDA, CpG, and AS03, we examined the ability of intranasally delivered CPAF, combined with these adjuvants alone and in combination, to protect female mice against intravaginal challenge with *Chlamydia muridarum*. While all adjuvant regimens were well-tolerated, only those containing CDA induced a memory T cell response, characterized by CD4 T cells producing IFNγ or IL-17A alone, or in combination with TNFα. Moreover, administration of CPAF with CDA alone was sufficient for protection, as evidenced by a significantly reduced chlamydial burden and faster clearance compared to controls. These findings demonstrate the potential for using STING agonists with CPAF in a prospective human vaccine against Chlamydia.

## MATERIALS & METHODS

### Mice

Female C57BL/6J (stock #: 000664) mice were purchased from The Jackson Laboratory (Bar Harbor, ME). Mice were given food and water ad libitum in an environmentally controlled, pathogen-free room with a cycle of 12 h of light and 12 h of darkness. Mice were age-matched and used between 8 and 12 weeks of age. All experiments were approved by the Institutional Animal Care and Use Committee at the University of North Carolina and the University of Chicago.

### CPAF antigen and adjuvants

An inactive variant of Chlamydia (*Chlamydia muridarum*) Protease Activity Factor (CPAF Cm) antigen was expressed using XpressCF® (Sutro Biopharma). The catalytic serine at position 491 was changed to alanine. The CPAF Cm clone was a gift from Jon Harris (Queensland University of Technology). After cell-free expression, the protein was purified by Ni affinity chromatography (Cytiva). CPAF Cm was admixed with CpG1826 (Invivogen) (10μg/dose) and/or STING agonist 2’3’-c-di-AM(PS)_2_ (Rp,RP) (Invivogen) (5μg/dose) in PBS or mixed at a 1:1 ratio with the squalene oil-in-water adjuvant AddaS03 (AS03) (Invivogen). As a comparator, CPAF was conjugated to 2Bxy-dopamine (Dopa) using SPAAC reaction^26^. In brief, 2.5x molar excess of 2Bxy-Dopa-DBCO was mixed with CPAF-N_3_ in PBS:PEG300:DMSO (7:2:1) and incubated at 35°C while shaking. Solution was washed with PBS through Amicon filter (10kDa) three times to remove DMSO. Reaction was verified with SDS-PAGE gel.

### SDS-PAGE Gel Electrophoresis

Proteins were prepared in 2x Laemmli sample buffer (Bio-Rad, Philadelphia, PA) added with 5% β-mercaptoethanol and denatured by heating at 95 °C for 5 min. The denatured protein samples (20 µl) were loaded onto 12% precasted polyacrylamide gels (Bio-Rad). For western blot analysis, SDS-PAGE-run samples were then transferred into the nitrocellulose membrane using Trans-Blot Turbo Transfer Packs (Bio-Rad). Then the membrane was blocked with PBS containing 0.5% Tween-20 (PBS-T) and 3% BSA, and incubated with anti-His tag antibodies (#MA1-21315, Invitrogen; diluted 1:5,000 in PBS-T) or murine anti-CPAF immune sera (pooled from mice immunized with CPAF protein; diluted 1:1000 in PBS-T) overnight at 4°C. The membrane was washed with PBS-T and incubated with Horseradish Peroxidase (HRP)-labeled goat anti-mouse IgG (#5220-0460, KPL SeraCare, Gaithersburg, MD) diluted 1:10,000 in PBS-T for 1 hour at room temperature. The membrane was again washed with PBS-T, revealed with Immobilon Western Chemiluminescent HRP Substrate (Millipore Sigma, Burlington, MA) and analyzed using Gel Doc imaging system (Bio-Rad). Blots and gels derived from the same or side-by-side experiments were processed together

### Analytical SEC

Analytical SEC experiments were performed with a Superdex 200 Increase 10/300GL SEC column (Cytiva). The flow rate was 1 ml/min in PBS running buffer. 200ul of a 1mg/ml protein sample was injected, and the chromatograms were monitored at A_280_.

### Mass spectrometry

The analysis was done on a Sciex X500B QTOF (AB Sciex, Redwood City, CA) coupled with Agilent’s 1290 UHPLC system (Agilent Technologies, Santa Clara, CA). LC-MS and LC-MS-MS approaches were taken to determine and confirmed the clip site of CPAF. A Waters (Waters, Milford, MA) BioResolve RP mAb Polyphenyl Column (450 Å, 2.7µm 2.1×50mm) column was used for intact protein analysis. Another Waters C18 column (Acquity Premier, Peptide CSH C18 130A 1.7uM, 2.1×150mm) was used for the peptide mapping analysis. The acquired MS data were analyzed with BioPharmaView (AB Sciex, Redwood City, CA) Software.

### Mucosal immunization

Mice were anesthetized intraperitoneally with 250 microliters of Nembutal (50 mg/mL) and then immunized intranasally on day 0 with 15μg adjuvanted CPAF or CPAF alone in a 12μl volume (6μl/nare). Mice were intranasally boosted 30 days later with the same vaccine formulation(s).

### ELISpot assay

Single-cell suspensions of murine splenocytes were prepared by passing cells through 70 μM cell strainers and ACK lysis buffer prior to resuspension in complete media. For analysis of IFNγ production, cells (1-2.5×10^5^ cells/well) were stimulated with a pool of 18-amino acid peptides overlapping by 15 amino acids (Sigma Aldrich) spanning the *Chlamydia muridarum* CPAF S491A antigen (final concentration of 2.5μg/ml)^27^ on PVDF-membrane plates (Millipore) coated with 5μg/ml anti-mouse IFNγ (AN18). After 18-20 hours of stimulation, IFNγ spot-forming cells were detected by staining membranes with anti-mouse IFNγ biotin (1μg/ml; R46A2) followed by streptavidin-alkaline phosphatase (1μg/ml) and developed with NBT/BCIP substrate solution (Thermo Fisher). Spots were enumerated on an AID ELISpot reader.

### Intracellular cytokine staining (ICS)

For analysis of intracellular cytokines, cells were stimulated as above in a round-bottom 96 well plate at 1×10^6^ cells/well in the presence of 2.5μg/ml CPAF pooled peptides (Synpeptide), media alone (negative control), or Cell Activation Cocktail (Biolegend; positive control) for 6 hours at 37°C with 5% CO_2_ in the presence of brefeldin A (Biolegend) and monensin (Biolegend). Cells were stained with a Zombie UV fixable viability kit (Biolegend) in PBS. Cells were washed and incubated with Mouse Fc Block (anti-CD16/CD32; BD Pharmingen) followed by staining for surface markers anti-Ly6G BV421 (1A8; Biolegend; dump channel), anti-CD45 Alexa Fluor 700 (30-F11; BioLegend), anti-CD3 FITC (17A2; BD Biosciences), anti-CD4 BV570 (RM4-5; BioLegend), anti-CD8 BUV395 (53-6.7; BD Biosciences), anti-CD44 BV480 (IM7; BD Horizon), and anti CD62L BV650 (MEL-14; Biolegend) in Brilliant Stain Buffer (BD Horizon). Following surface staining, cells were fixed and permeabilized using Cytofix/Cytoperm Fixation/Permeabilization Kit (BD Biosciences). Cells were then stained for intracellular cytokines anti-TNFα BV711 (MP6-XT22; BioLegend), anti-IFNγ APC Fire-750 (XMG1.2; BioLegend), anti-IL17A PE-Cy7 (TC11-18H10.1; BioLegend), and anti-IL-5 PE (TRFK5; Biolegend) using Cytofix/Cytoperm Fixation/Permeabilization Kit (BD Biosciences). Samples were washed and resuspended in PBS+1%FBS prior to acquisition on a Cytek Aurora spectral cytometer, and data were analyzed with FlowJo version 10 software.

### Antibody ELISA assay

Serum from vaccinated C57BL/6 mice were assayed for CPAF-specific IgG, IgG1, IgG2b and IgG2c antibody responses. ELISA plates were coated overnight at 4°C with 10μg/ml recombinant MBP-CPAF (from John Harris, Queensland University of Technology) diluted in 0.5M NaHCO3. Plates were washed with PBS-Tween. After blocking with 2% BSA PBS-Tween for 1 hour at 37°C, samples were serially diluted and incubated for 1 hour at 37°C. Internal controls were generated using reference serum. Plates were washed with PBS-Tween. A goat anti-mouse horseradish peroxidase-conjugated secondary antibody was added (Southern Biotech) and incubated for 1 hour at 37°C. Goat anti-mouse IgG was added at a 1:4,000 dilution and IgG1, IgG2b, and IgG2c were added at a 1:500 dilution. After washing, the plates were developed using TMB substrate solution (ThermoScientific) for a maximum of 20 minutes. The reaction was stopped using 0.5M H2SO4 and OD was read at 450nm. The cut-off for detecting a specific antibody response was defined by a value greater than the mean+3 standard deviations of the non-specific antibody values.

### Cytokine ELISA assay

Mouse lungs were harvested using T-PER Tissue Protein Extraction Reagent (Thermo Fisher, # 78510), supplemented with a protease inhibitor. The supernatant of lung homogenates was collected and used for ELISAs. Cytokine levels of TNF-α, IL-6, and IFN-γ were measured using ELISA Max kits from BioLegend, per the manufacturer’s instructions.

### Strains, cell lines, and culture conditions

Plaque-purified *C. muridarum* Nigg strain CM006^28^ was propagated in mycoplasma-free L929 cells^29^ and titrated by inclusion-forming units^30^ using a fluorescently tagged anti-chlamydial lipopolysaccharide monoclonal antibody (Bio-Rad).

### Chlamydia infection quantification

Female mice at least 8 weeks old were s.c. injected with 2.5 mg medroxyprogesterone (Depo-Provera; Upjohn) 5-7 days prior to infection to induce a state of anestrous. Mice were intravaginally inoculated with 1×10^5^ inclusion forming units (IFU) CM006 diluted in 30μl sucrose-sodium phosphate-glutamic acid buffer 28 days post booster immunization. Mice were monitored for cervicovaginal shedding via endocervical swabs^30^, and IFUs were calculated, as described previously^31^. Animal welfare was monitored daily.

### Lung histopathology

Lungs were perfused with PBS (2 mM EDTA) followed by 10% formalin. They were then inflated through the trachea using 10% formalin and fixed for 48 hours before paraffin processing. A pathologist blinded to the study design analyzed H&E-stained slides for gross pathology for inflammation and hemorrhaging. Inflammation scores were assigned as ‘0’ for none/negligible inflammation or slightly increased inflammatory cell presence only associated with areas of hemorrhage; or as ‘1’ for concentrated, prominent inflammation independent of areas with hemorrhage. Hemorrhage was classified as ‘none’; or if present, as ‘focal’ for hemorrhage seen in only one area of tissue sample or as ‘multi-focal’ for hemorrhage seen in ≥2 areas of the sample. For multi-focal hemorrhage, further classification was designated as ‘minimal’ for hemorrhage involving <25% of sample area; ‘moderate’ for 25-50% of sample area; and ‘diffuse’ for >50% of sample area.

### Hydrosalpinx and oviduct histopathology

Genital tract gross pathology, including hydrosalpinx development, was examined and recorded at sacrifice on day 42 post-challenge. The genital tract was collected in its entirety with careful removal of adipose as needed. Samples were placed flat between two sponges in a histology tissue cassette and placed in 10% neutral buffered formalin for at least 72 hours before routine processing to paraffin blocks. Tissue blocks were sectioned to 5 µm onto charged slides and stained with hematoxylin and eosin.

Histological samples were evaluated in a masked fashion by a board-certified veterinary pathologist (RSS). Within a single animal, each side was scored independently. An overall assessment of oviduct dilation was reported for each uterine horn. Samples were scored on a 0-5 scale: 0=no finding; 1=minimal finding; 2=mild finding; 3=moderate finding; 4=marked finding; 5=severe finding.

### Statistical Analysis

Differences between the means of experimental groups after infection were calculated using two-way repeated measures (RM) ANOVA. Significant differences in ELISA and ELISpot data were determined by one-way ANOVA. Significant differences in ICS responses were determined by the Wilcoxon Rank Sum Test. Statistical differences in oviduct histopathology were determined by the Kruskal-Wallis test. Prism software (GraphPad) was utilized for statistical analyses, and values of p ≤ 0.05 were considered significant.

## RESULTS

### Expression of inactivated C. muridarum CPAF using cell-free protein synthesis

The S491A mutation in the tail-specific protease domain renders the CPAF protease inactive and incapable of natural auto-processing and proteolysis^32, 33^. However, cell-free expression yielded full-length CPAF and a clipped form consisting of N- and C-terminal fragments (Fig. 1A). Mass spectrometry revealed the clip site to be at K232 (Supp. Fig. 1, Supp. Table 1), three amino acids from the native auto-processing site. The K232Q mutation produced a full-length Cm CPAF protein (Fig. 1B). Both Cm CPAF variants were applied to an analytical sizing column and eluted as a single Gaussian peak (Fig. 1C). The full-length wild-type and K232Q variant had the same retention time demonstrating similar folding and dimerization. The clipped form was used for vaccine studies.

**Figure 1.**
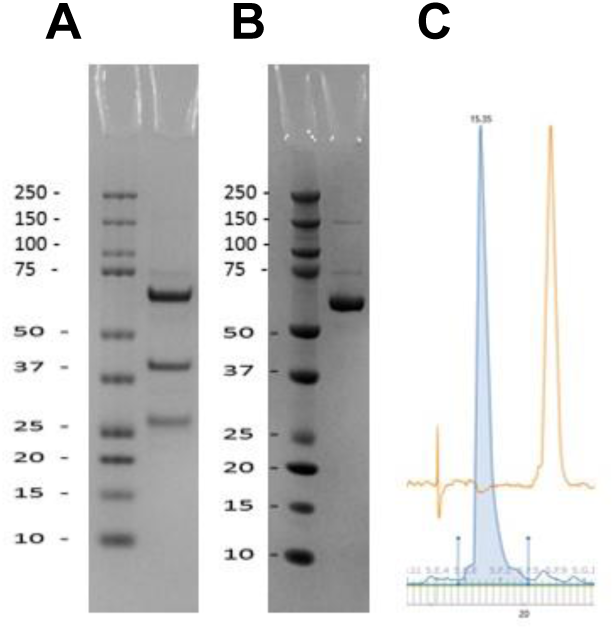
Recombinant *Cm* CPAF S491A expression in a cell-free protein synthesis platform. (A) SDS-PAGE showing full-length inactive CPAF and N-terminal and C-terminal protein fragments. (B) SDS-PAGE showing unclipped full-length K232Q inactive CPAF. (C) Analytical size-exclusion chromatography for both CPAF variants.

### Intranasal immunization with adjuvanted CPAF does not induce lung pathology

Female mice were immunized intranasally with CPAF formulated with different adjuvant combinations and protein/vehicle controls. Lungs were harvested at 3-hour, 24-hour, and 1-week time points post-immunization for analysis of inflammatory cytokines and pathology (Fig. 2A-C). Lung homogenate levels of IFNγ, IL-6, and TNFα after immunization were not significantly different than PBS vehicle controls across all time points. Further, lung histology did not show signs of inflammation or differences in multi-focal hemorrhaging between the PBS vehicle group and adjuvanted formulations. No signs of necrosis or scare tissue were observed. (Fig. 2D-F)

**Figure 2.**
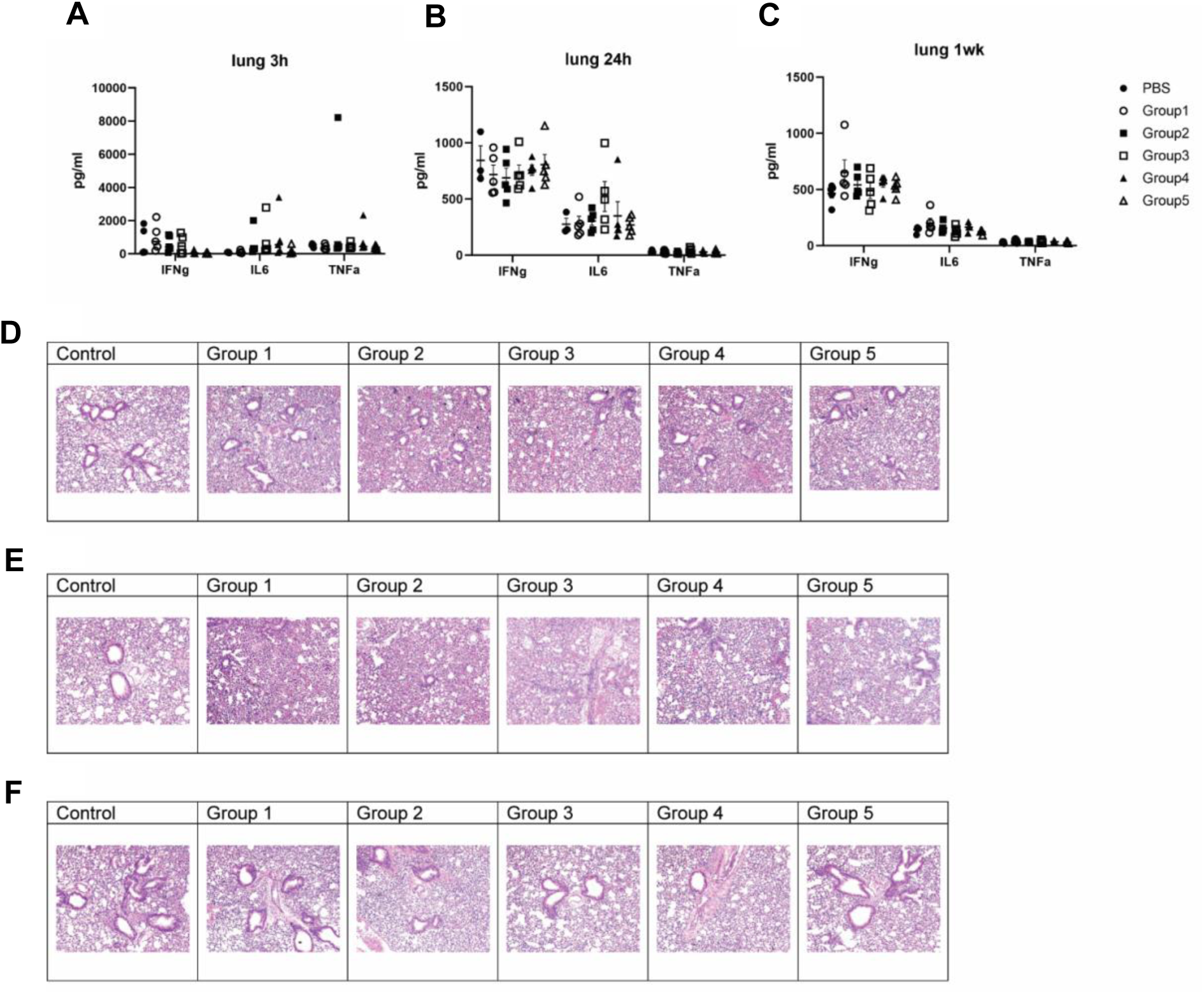
Intranasal administration of adjuvanted CPAF does not induce inflammation or gross pathology in the lung. Levels of IFNγ, IL-6, and TNFα in the lung 3 hours (A), 24 hours (B), and 1 week (C) post immunization. H&E staining of lungs from immunized mice (n=5/group) after 3 h hours (D), 24 hours (E), and 1 week (F) post-immunization. Group 1: CPAF only, Group 2: CPAF + CpG + CDA, Group 3: CPAF + CpG + CDA + AS03, Group 4: CpG + CDA + AS03, Group 5: CPAF-2Bxy-dopa (TLR7/8 agonist).

### Intranasal immunization with CPAF plus CDA induces a memory CD4 T cell response characterized by the production of IL-17A or IFNγ

Female C57BL/6 mice immunized intranasally and boosted with a second dose thirty days later were assessed for systemic antibody and T cell responses ten days post-boost (Fig. 3A). Most mice immunized with CPAF alone, CPAF plus CDA, or CPAF plus CpG/CDA/AS03 had low to undetectable serum antibody titers with no significant differences between the groups (Fig. 3B). Th1-biased IgG2b and IgG2c were detected in a few individual mice, while IgG1 was not detected. In contrast, mice immunized with CPAF plus CDA alone or in combination with CpG and/or AS03 had elevated levels of CPAF-specific IFNγ-producing T cells (mean = 562-676 SFUs) (Fig. 3C), which were not consistently detected in mice immunized with CPAF alone or adjuvanted with CpG and/or AS03. We did not observe any synergistic effect when CpG and/or AS03 were included with CDA. The memory (CD44^hi^ CD62L^-^) CPAF-specific T cell response after CPAF + CDA immunization was almost entirely CD4-biased and contained a mixture of CD4 T cells producing either IL-17A or IFNγ with very few IL-17A/IFNγ co-producing cells (Supp Fig.2, Fig. 3D-E). Nearly 4% of memory CD4 T cells produced TNFα alone or in combination with IL-17A or IFNγ, while TNFα was the only cytokine significantly detectable over background from CD8 T cells.

**Figure 3.**
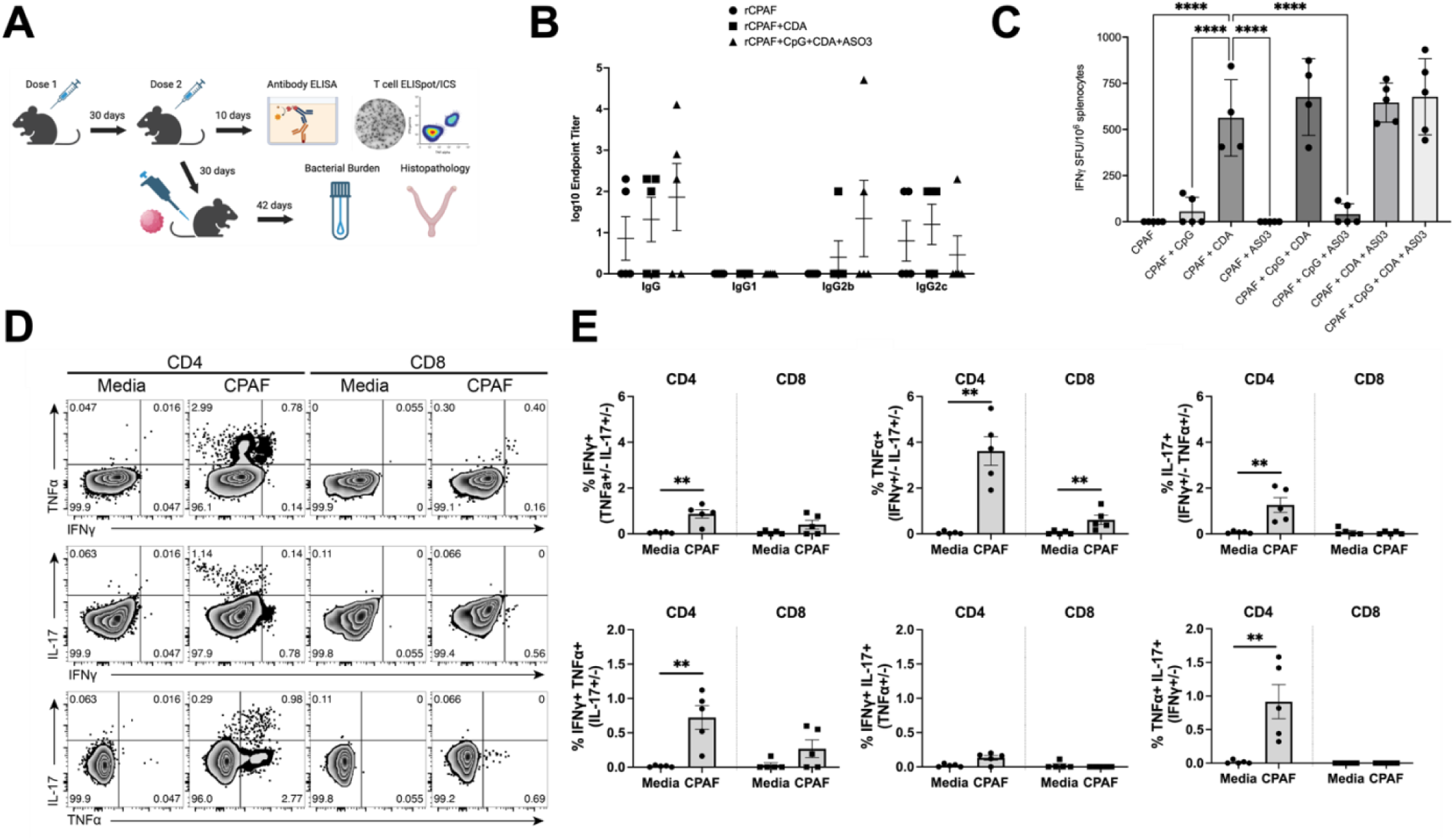
Cellular and humoral responses after intranasal CPAF immunization. (A) Schematic for immunogenicity and challenge experiments. (B) Antibody titers in mice immunized with CPAF plus CDA alone, triple adjuvant combination, or antigen alone. p=NS for all comparisons by one-way ANOVA (n=5/group). (C) IFNγ ELISpot responses in immunized mice for all adjuvant iterations (n=4-5/group). ****p<0.0001 by one-way ANOVA (D) Representative plots for CPAF-specific cytokine responses in CPAF + CDA immunized mice by ICS (n=5). (E) Frequency of CPAF-specific mono- and poly-functional memory (CD44^hi^ CD62L^-^) CD4 and CD8 T cell responses in CPAF + CDA immunized mice. **p<0.01 by Wilcoxon rank-sum test.

### Intranasal immunization with CPAF plus CDA reduces cervical burden in female mice

Female mice intranasally immunized with CPAF plus CDA alone or combined with CpG and/or AS03 had significant reductions in cervical chlamydial burden after intravaginal challenge compared to PBS controls (Fig. 4A), with CPAF plus CDA immunization resulting in a 1.5 log reduction in cervical burden. We observed no synergistic protection when combining CDA with CpG and/or AS03. The only regimen that significantly reduced bacterial burden and the frequency of oviduct hydrosalpinx, compared to PBS controls, was CPAF plus CDA (Fig. 4A-C). In contrast, the unadjuvanted controls and formulations that included AS03 without CDA exhibited high frequencies of oviduct hydrosalpinx (Fig. 4B).

**Figure 4.**
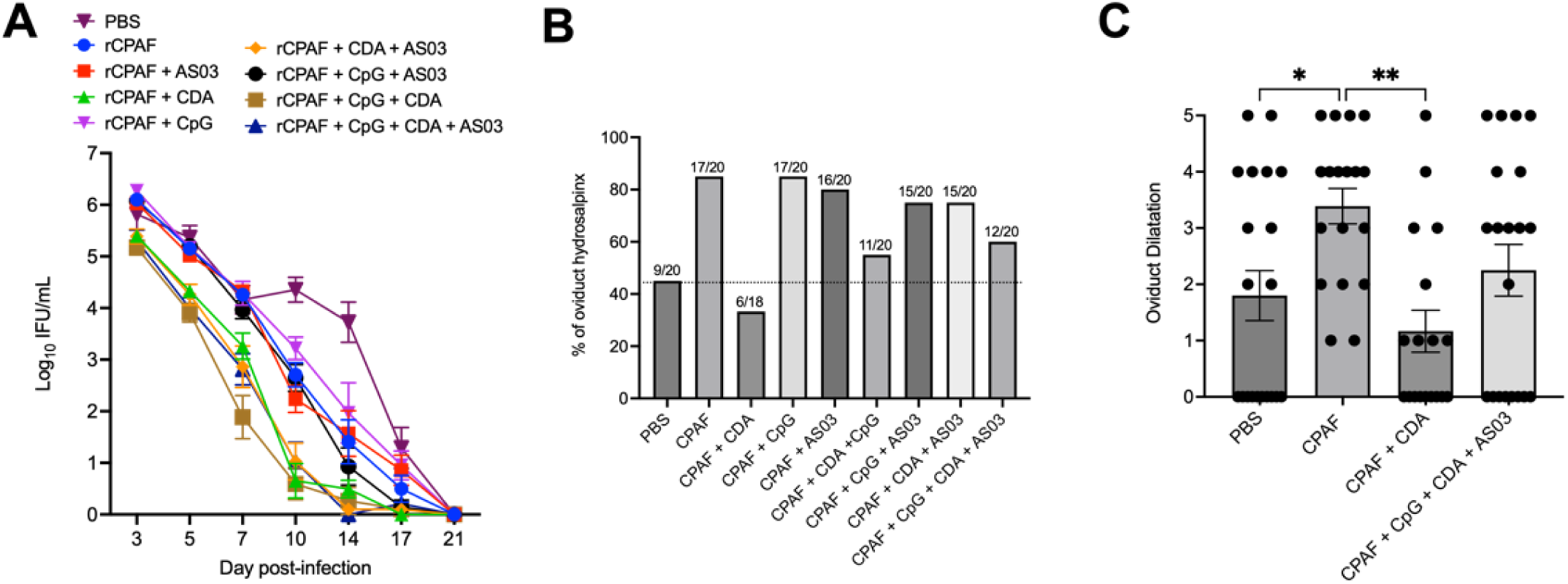
CPAF plus CDA intranasal immunization reduces burden in intravaginally challenged mice and does not enhance oviduct pathology. (A) Course of *C. muridarum* challenge infection in immunized mice and controls (n=9-10 mice/group). rCPAF + CDA vs. PBS (−1.51 log, ****p<0.0001), rCPAF + CDA vs. rCPAF (−0.85 log, ****p<0.0001), rCPAF + CDA vs. rCPAF + CpG (−1.10 log, ****p<0.0001), rCPAF + CDA vs. rCPAF + AS03 (−0.84 log, ****p<0.0001), rCPAF + CDA vs. rCPAF + CpG + AS03 (−0.69 log, ****p<0.0001), p=NS for rCPAF + CDA vs. all vaccine combinations incorporating CDA. Significance determined by two-way repeated-measures ANOVA. (B) Frequency of oviduct hydrosalpinx after challenge of mice immunized with CPAF plus CDA alone, triple adjuvant combination, CPAF alone, or PBS. Dotted line set to oviduct hydrosalpinx frequency in PBS controls. (C) Oviduct dilatation scores of immunized mice and controls. Significance determined by Kruskal-Wallis test. *p<0.05, **p<0.01

## DISCUSSION

STING pathway agonists have demonstrated an excellent safety profile in mice and humans^16, 17^ and have elicited strong T cell immunogenicity and protection against bacterial pathogens like *Mycobacterium* tuberculosis^34, 35^ and *Bordetella* pertussis^36, 37^. However, their efficacy has not been tested against Chlamydia infection. Thus, we hypothesized that intranasal immunization with the immunodominant secreted antigen CPAF combined with CDA would induce a memory CD4 T cell response and protect female mice against genital Chlamydia infection.

We expressed full-length inactive CPAF in a cell-free protein synthesis platform amenable to scale-up^38^. While clipped form was used in these studies, we were recently able to express a full-length form that provides a consistent product for manufacture and will be the focus of future analyses. Mass spectrometry revealed *in vitro* clipping at the Lysine232 residue that is three amino acids upstream of the natural auto-processing cleavage site^32, 33^. Mutation of this residue to glutamine prevented clipping to yield a full-length product. These data suggest the loop region of CPAF is especially prone to proteolysis, perhaps both by auto-processing and exogenous proteolytic activity from bacterial cell lysate.

Intranasal delivery of CPAF with CDA, CpG, and/or AS03 did not induce any detectable damaging inflammatory responses in the lungs after intranasal immunization, aligning with the known safety records of these adjuvants in mice and humans. We did not observe any significant increases in the levels of IFNγ, IL-6, or TNFα in the lungs compared to PBS controls, nor differences in multi-focal hemorrhaging compared to unadjuvanted controls. These results are promising for future investigations of intranasal vaccines using these adjuvants against other pathogens.

Prime-boost immunization induced low to undetectable frequencies of CPAF-specific IgG. However, the few detectable responses were characterized by Th1-biased IgG2b and IgG2c consistent with the antibody profiles of mice immunized intranasally with CPAF adjuvanted with CpG or IL-12^14, 15^. Therefore, CPAF-specific antibodies are unlikely to play a significant role against this secreted antigen. Recent data showed that B-cells were completely dispensable for protection elicited with chimp adenovirus (ChAD)-vectored CPAF^27^, and B-cell deficient mice immunized with recombinant CPAF did not exhibit an altered course of clearance^39^. However, it is possible that Fc-mediated antigen presentation could be enhanced by CPAF-specific IgG, resulting in boosted CD4 T cell responses^40^.

We found that the inclusion of CDA in the adjuvant formulation was necessary for T-cell immunogenicity. Adding CpG and/or AS03 to CDA did not have a synergistic effect on the frequency of IFNγ producing T cells, unlike in some other vaccine models^41^. CPAF-specific T cells were characterized by a CD4-dominant response, consistent with other studies using protein immunization^42^. We did not observe a significant CPAF-specific CD8 T cell response, which contrasts sharply with the dominant CD8 T cell response elicited with ChAd.CPAF^27^. This difference can be explained by the fact that exogenous recombinant proteins are easily endocytosed by professional antigen-presenting cells for MHC class II presentation to CD4 T cells, while viral vectors predominately release antigens into the infected cell cytosol for presentation on MHC class I to CD8s^43^.

The CD4 response demonstrated a mixed profile of IFNγ ± TNFα and IL-17A ± TNFα producing cells, like those achieved in mice with the subcutaneously delivered MOMP-based immunogen CTH522 adjuvanted with CAF01^44^. The immunophenotypic T cell expression of IFNγ ± TNFα is routinely measured in Chlamydia studies, while detecting IL-17A responses is less common^45^. It is unclear if there is a need for de facto or ex-Th17 cells in vaccine-elicited protection against Chlamydia genital infection, as observed for other bacterial pathogens like *Klebsiella spp*. and *Mycobacterium* tuberculosis^46, 47^. We also observed a low frequency of TNFα single-positive CD8 T cells. These monofunctional cells are shown to be dispensable for protection^27^.

The vaccine-elicited reductions in chlamydial burden after genital challenge correlated with our T cell immunogenicity data. Mice vaccinated with regimens including CDA exhibited similar clearance kinetics and achieved a 1.5-log reduction in bacterial load compared to PBS controls. These results further confirmed that CDA alone was sufficient for protection. Despite significant reductions in burden, we were not able to significantly prevent the development of hydrosalpinx with our CPAF + CDA regimen. However, a trend for reduced gross hydrosalpinx and oviduct dilatation compared to PBS controls was observed, and this regimen did not result in the levels of oviduct dilatation elicited with unadjuvanted CPAF or non-protective regimens incorporating AS03 with CDA. While beyond the scope of these analyses, it is possible that these non-protective adjuvant formulations lead to tissue-damaging immune responses in the oviduct.

These studies further highlight the effectiveness of STING agonists in providing protection against infectious pathogens, and their documented clinical safety and immunogenicity in humans make them an attractive adjuvant. Potential improvements to improve this platform include incorporating additional immunogens or doses, utilizing more effective delivery vehicles, and exploring covalent conjugation of agonists. Future studies will focus on enhancing the efficacy of this recombinant CPAF vaccine, with the goal of advancing to human clinical trials.

## ACKNOWLEDGEMENTS

The UNC Flow Cytometry Core Facility (RRID:SCR_019170) is supported in part by P30 CA016086 Cancer Center Core Support Grant to the UNC Lineberger Comprehensive Cancer Center.

