## Supplemental Figures 1 and 2 and Table 1 for "Intranasal immunization with CPAF combined with cyclic-di-AMP induces a memory CD4 T cell response and reduces bacterial burden following intravaginal infection with *Chlamydia muridarum*"

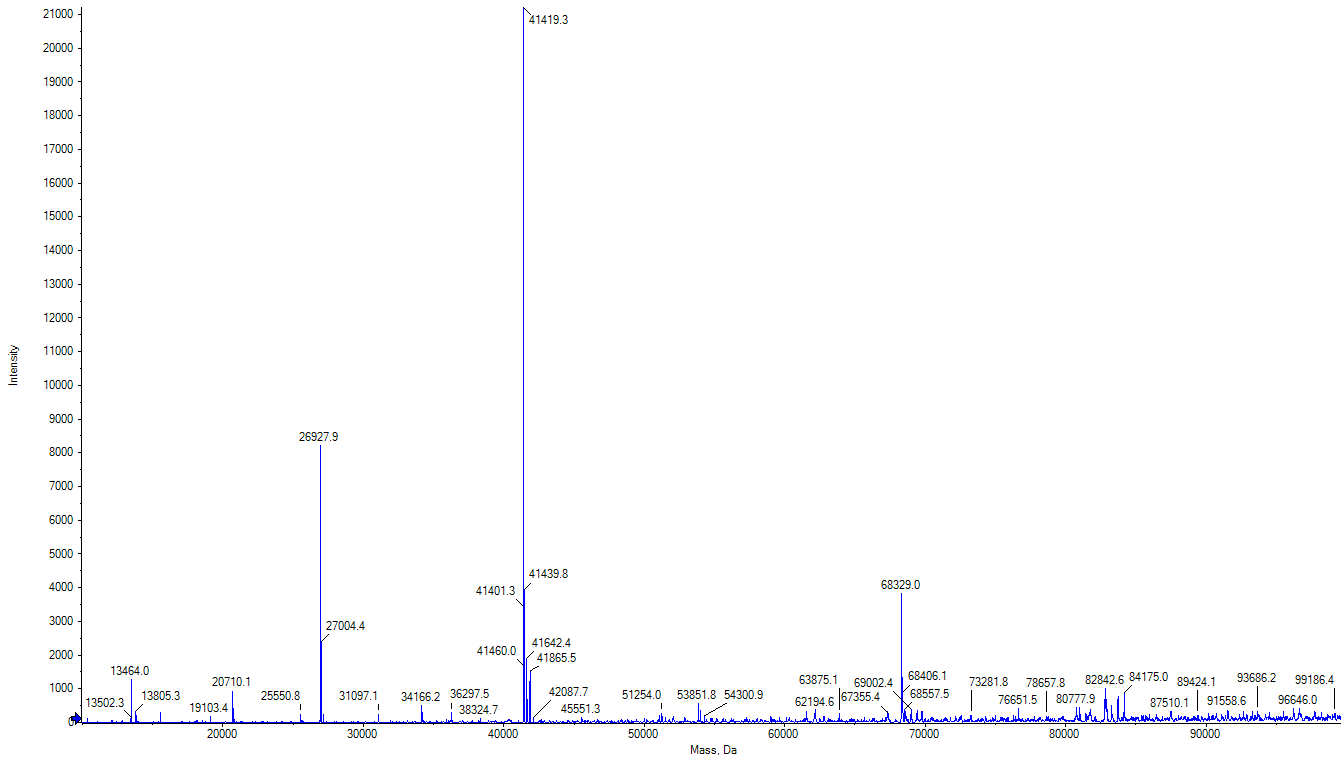

**3**

**2**

**1**

**Supplementary Figure 1. Deconvoluted mass spectrum of intact protein analysis by LC-MS showed the three major forms of S491A CPAF: 1) the N-terminal clipped fragment; 2) the C-terminal fragment; and 3) the full length protein. The X-axis indicates the mass spectrometry signal intensity, the Y-axis indicates the mass of the protein or protein fragments in Dalton.**

| **Fragment** | **Position to N-terminal** | **Theoretical Mass Dalton** | **Observed mass Dalton** | **ΔDalton** |
| --- | --- | --- | --- | --- |
| N-terminal | 1-244 | 26928.3 | 26927.3 | -1.0 |
| C-Terminal | 245-613 | 41420.0 | 41419.3 | -0.7 |
| Full Length | 1-613 | 68330.3 | 68329.0 | -1.3 |

**Supplementary Table 1. The molecular weight in mass Dalton of the three forms of S491A measured by LC-MS. The result showed that the clipping site was between position 244 and 245 from the N-terminal. The mass difference relative to the theoretical mass was within 1 Dalton. This clipping site was further confirmed by LC-MS-MS.**

**Supplementary Figure 2. Flow cytometric gating strategy for detection of CPAF-specific memory CD4 and CD8 T cell responses**
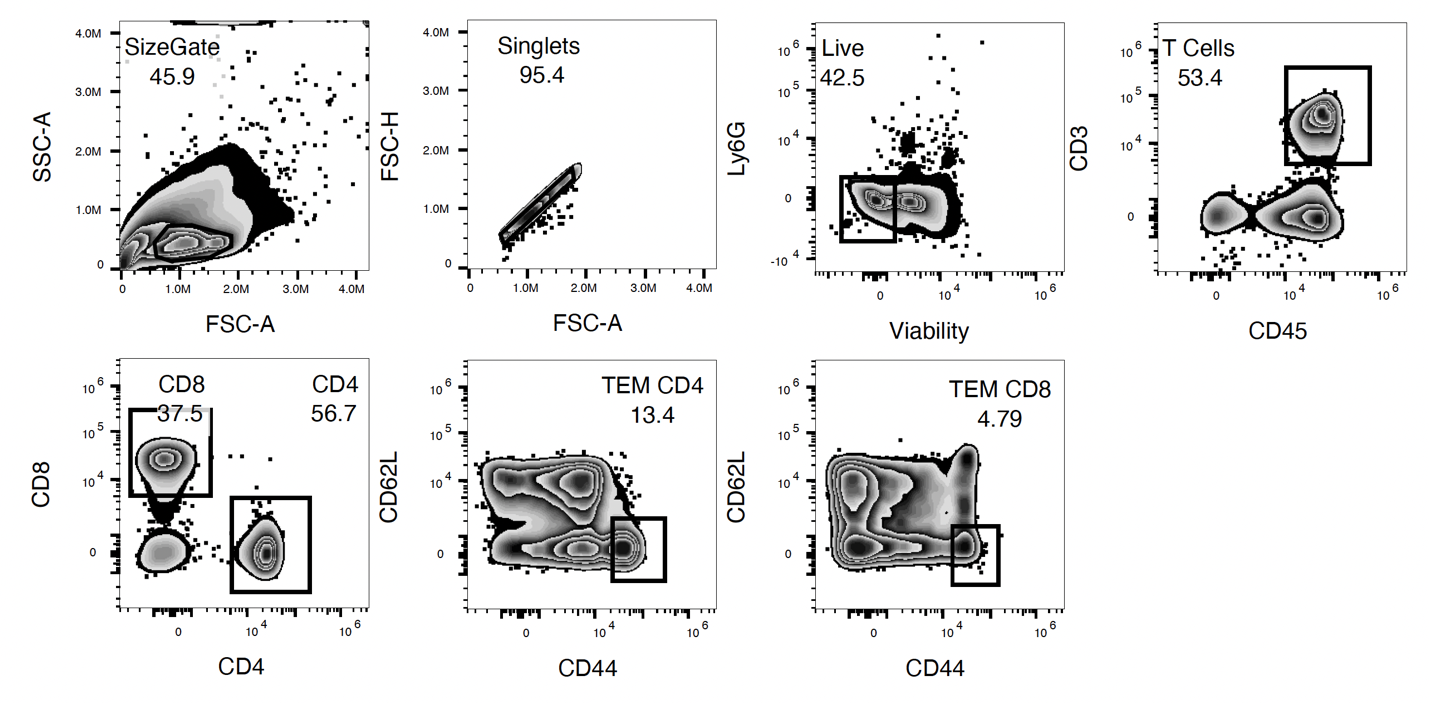
**.**
